# Potato dihaploids uncover diverse alleles to facilitate diploid potato breeding

**DOI:** 10.1101/2025.08.06.668948

**Authors:** Sapphire Coronejo, Brieanne Vaillancourt, John P. Hamilton, Xiaoxi Meng, Kathrine Mailloux, Grace Christensen, Jessica Huege, Kathleen M. Shaw, Husain I. Agha, Kristen Brown-Donovan, James S. Busse, Andy Hamernik, Maria V. Caraza-Harter, Lucas Heroux, Hemant Balasaheb Kardile, Ericka Knoeck, Peyton L. Sorensen, Diana Spencer, Solomon Yilma, Paul C. Bethke, David Douches, Joshua Parsons, Vidyasagar Sathuvalli, Ek Han Tan, Jeffrey B. Endelman, C. Robin Buell, Laura M. Shannon

## Abstract

Commercial potato in North America is a clonal autotetraploid crop, which complicates breeding. Efforts are underway to convert potato to a diploid inbred-hybrid crop that is amenable to additional breeding strategies and allows breeders to more quickly respond to demands for crop improvement. With the goal of preserving haplotypes developed over the last 200 years of selection, diploid potato breeding in the US started with the creation of diploids from tetraploid commercial varieties and advanced breeding lines through prickle pollination. This is an effective but slow method which presents a barrier to entry for individual breeding programs. Therefore, we developed 97 publicly available dihaploids (diploids from prickle pollination of tetraploids) as a resource for diploid breeding in the U.S. These clones contain the majority of alleles in the US breeding population for three market classes: chips, russets, and fresh market reds. To facilitate genomic informed breeding, all clones have been resequenced using short read sequencing technology, and we have developed *de novo* assemblies based on PacBio HiFi long reads for 20 individuals. As an illustration of how these data will be used in breeding, we explored the maturity locus (*StCDF1*) and identified 15 different alleles. The majority of dihaploids were heterozygous for early and late alleles, resulting in intermediate maturity. Beyond informing breeding, this data facilitates investigations into potato genomics. The dihaploid population is both highly heterozygous and incredibly diverse on a population level. In particular, there is extensive structural diversity, including copy number variation, segregating within the population. This contrasts with a relatively low genome-wide historical recombination rate (*ρ*). Taken together, these findings indicate that potato is highly diverse, with much of that diversity found within long linkage blocks.

## Introduction

Potato is the most widely grown vegetable crop in the world. It is central to cuisines and cultures across the globe, in part, due to its high protein, fiber, potassium, vitamin C, and antioxidant content (Beals, 2019; Navarre et al., 2019). Changes in the environment and consumer preferences necessitate new potato varieties, yet development of new varieties is challenging due to the clonal and autopolyploid nature of cultivated potato. Clonal propagation is slow, resulting in considerable time to market for new varieties, and risks transmission of disease agents. Autopolyploidy reduces the effect of selection (Monnahan & Brandvain, 2020), permits the accumulation of deleterious alleles (Booker & Schrider, 2025), and makes inbreeding and introgression difficult (Gallias, 2003). Finally, potatoes are a vegetable with many quality attributes; appearance, processing, and storage traits are all essential to marketability and must be balanced with agronomic traits during selection. Taken together, clonal growth, autopolyploidy, and the numerous diverse breeding targets makes progress in potato breeding inherently slow.

One way to improve the effectiveness of potato breeding is to reinvent potato as a diploid inbred-hybrid crop based on true potato seed (Lindhout et al., 2011; Jansky et al., 2016; Zhang et al., 2021; Bradshaw, 2022; de Vries et al., 2023). The numerous benefits of this system include the ability to take advantage of commonly used breeding techniques such as inbreeding, trait fixation, heterosis, and introgression, as well as simplifying seed increase, transportation, storage, and disease control. Although there were concerns that tetraploid-derived tubers tended to be larger than diploid-derived tubers, diploid tubers can compete with tetraploid tubers in terms of size (Alsahlany et al. 2021) and yield (Marand et al. 2019). Early efforts at diploid breeding have fixed traits in diploid potato germplasm (Song & Endelman, 2023) and shown evidence of heterosis (Zhang et al., 2021).

There are multiple possible sources of diploid potato breeding germplasm. Many South American landraces and commercial varieties are diploid as is much of the international genebank collection (Anglin et al., 2024). These diploid clones are currently being used as the basis of breeding programs through a process of re-improvement (Zhang et al., 2021). Wild potatoes are also primarily diploid and play large roles in the pedigrees of historically developed diploids available through the Agriculture-AgriFood Canada and USDA genebank (Achakkagari et al., 2022). While these approaches are promising, they inevitably result in the loss of alleles developed over the last 200 years of breeding, which are essential to the potato processing industry in the United States (U.S.). In the interest of maintaining these alleles from tetraploid cultivated potatoes, we developed dihaploid (diploid) populations from existing tetraploids, both commercial cultivars and advanced breeding lines. This was achieved through pollination with haploid inducers to generate maternal dihaploids (Busse et al., 2021; Hougas and Peloquin 1957; Hougas and Peloquin 1958; Hougas et al. 1958). The resulting diploid offspring have half the number of chromosomes of the tetraploid parent and minimal contribution from the inducer (Amundson et al., 2021; Pham et al. 2019).

Haploid induction through prickle pollination, as described above, has low efficiency (Busse et al., 2021), and generating dihaploids as a starting population for a diploid breeding program represents a serious time commitment and barrier to entry. Furthermore, a breeding program with a small founder population runs the risk of fixing deleterious alleles (Agha et al., 2023), which are present at a high frequency in the potato genome (Zhang et al., 2019; Zhang et al., 2021; Hoopes et al. 2022; Wu et al., 2023). Therefore, we aimed to generate a population of dihaploids that encompasses the diversity in U.S. tetraploid potatoes to provide a foundation for diploid-based breeding programs; all 97 individuals in this set, the Potato 2.0 Dihaploid Panel (P2DP), are available for research and breeding via the U.S. Potato Genebank in Sturgeon Bay, Wisconsin. With the goal of supporting genomic-informed breeding in diploid potato, we sequenced all 97 lines using Illumina short reads and generated PacBio HiFi long read *de novo* assemblies for 20 individuals. We used these data to assess allelic diversity, structural variation, and recombination frequencies in U.S. cultivated potato germplasm, revealing key information that can be used in genome-enabled breeding programs.

In addition to serving as a resource for breeders, this data set provides a unique opportunity to better understand the genome of tetraploid potato and a tool for genetics. Autotetraploid genomics are complex and dihaploids provide the opportunity to study tetraploid alleles using diploid technologies (Manrique Carpintero et al., 2018; Akai et al., 2023; Endelman et al., 2024).

## Results

### The Potato 2.0 Dihaploid Panel

The P2DP was generated from 58 cultivated tetraploids representing the (1) Chip processing market class (Chips), (2) Red skinned fresh market class (Reds), (3) Russet market class (Russets) and (4) Other clones (Other) which include specialty potatoes with yellow or purple flesh as well as heirloom varieties (Supplemental Table 1). Whole-genome resequencing generated 106 to 251 million reads per dihaploid, with alignment rates to the doubled monoploid reference genome DM 1-3 516 R44 v6.1 (hereafter DM) ranging from 90.6% to 99% with an average alignment rate >98% for each market class (Table 1, Supplemental Table 1). Of the 100 dihaploids sequenced, 97 were confirmed as diploid, one as triploid, and two as tetraploid (Supplemental Table 2); the 97 dihaploids, having stable and confirmed ploidy, were selected for further analyses.

**Table 1.** Summary of sequence data and mapping statistics.

| Population | Number of clones | Total Reads<sup>a</sup> | GC content (%)<sup>a</sup> | Mapped Reads<sup>a</sup> | Mapped Reads (%)<sup>a</sup> |
| --- | --- | --- | --- | --- | --- |
| All | 97 | 182,209,334 | 35.0 | 179,750,094 | 98.7 |
| Chips | 40 | 196,812,639 | 35.0 | 193,767,833 | 98.5 |
| Reds | 20 | 175,482,931 | 35.0 | 173,355,472 | 98.8 |
| Russets | 29 | 165,534,706 | 35.0 | 163,526,492 | 98.8 |
| Others | 8 | 186,454,341 | 34.9 | 184,458,509 | 98.9 |

<sup>a</sup> average

To understand haplotype structure at the chromosome-level, 20 diverse dihaploids were sequenced using PacBio HiFi technology, assembled into haplotype-phased assemblies, and scaffolded into chromosomes using reference-based scaffolding (Supplemental Table 3, Supplemental Figure 1). The 20 assemblies had an average haploid genome assembly length of 795 Mb and average N50 scaffold length of 64 Mb. Overall, the two haplotypes within each dihaploid were comparable in size, suggesting sufficient sequence diversity between the haplotypes (Supplemental Figure 2, Supplemental Table 3). The quality of the assemblies was evidenced by an average per haplotype-resolved assembly of 98.2% complete Benchmarking Universal Single Copy Orthologs (BUSCO), of which, 95.2% were complete and single copy vs. 2.9% complete and duplicated (Supplemental Figure 3, Supplemental Table 3).

Each genome was repeat masked with the majority of repetitive sequence being retroelements (Supplemental Table 4). Each dihaploid was then annotated for protein-coding genes using a uniform annotation pipeline that included full-length cDNA sequences from leaf and tuber resulting in an average of 87,287 working gene models per dihaploid, of which, 68,742 were high confidence gene models encoded by 62,634 high confidence genes (Supplemental Table 5). Representation of complete BUSCO orthologs in the high confidence representative gene set was 92.1% to 93.6% across the dihaploids with the majority being duplicated due to the presence of two haplotypes within each dihaploid assembly (Supplemental Table 6).

### Dihaploids exhibit high individual and population level diversity

We measured diversity at the individual, market class, and population level (Figure 1). First, for each dihaploid, we calculated observed heterozygosity (*H_O_*) using 36.2 million phased SNPs (Figure 1B and Table 2). *H_O_* differed significantly between market classes (*p* = 5.49x10^-4^). Overall, Chips had the highest mean *H_O_* at 0.81% while Russets had the lowest mean *H_O_* at 0.75% (Figure 1B) (*p* = 5.24x10^-4^, Dunn’s test). In general, Chips and Others showed similar levels of *H_O_* as did Reds and Russets, but pairwise comparisons between these two groups were significantly different (Chips vs. Reds: *p* = 9.51x10^-3^; Chips vs. Russets: *p* = 5.24x10^-4^; Reds vs. Others: *p* = 3.14x10^-2^; Russets vs. Others: *p* = 1.74x10^-2^).

**Figure 1.**
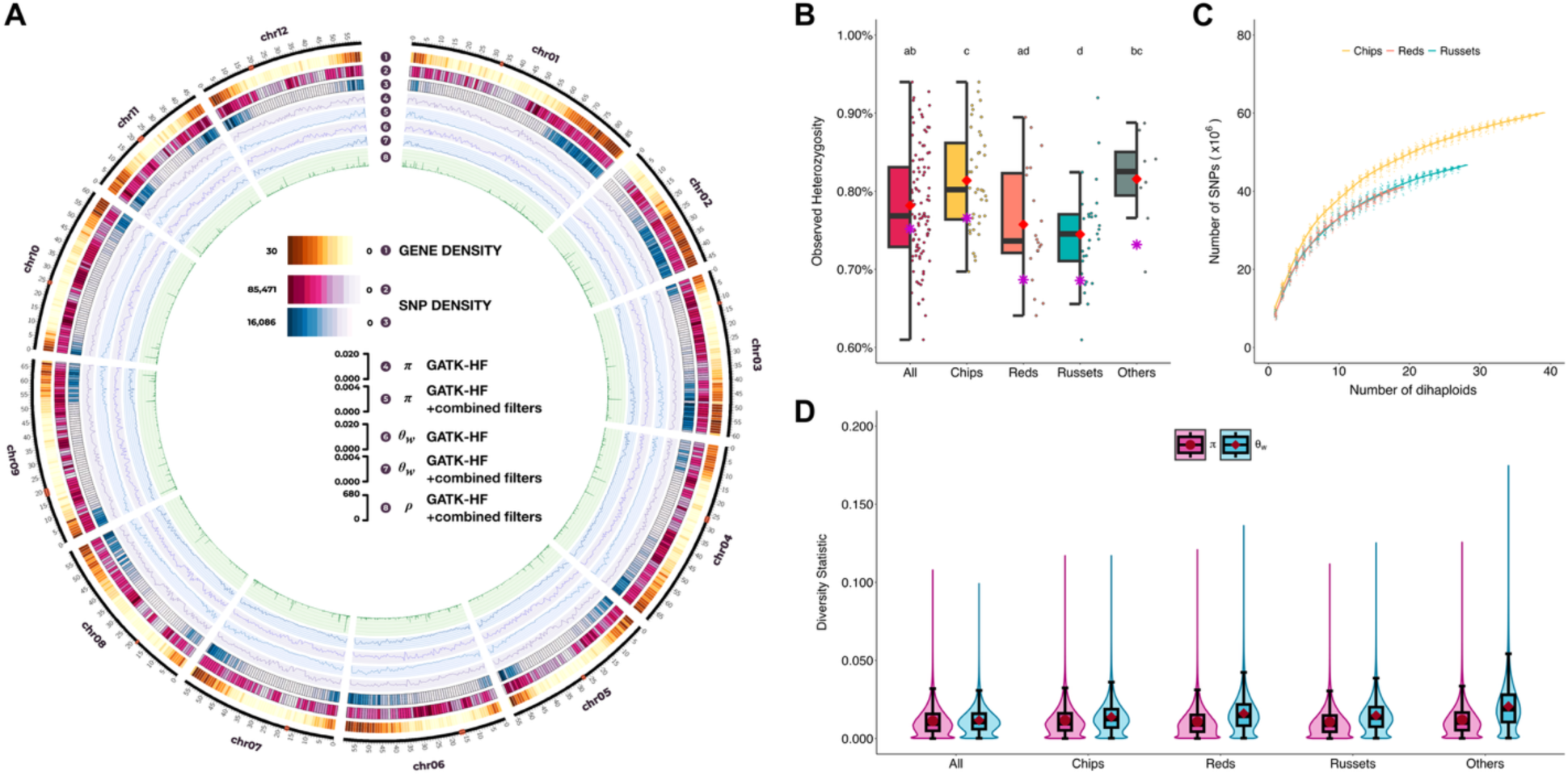
Genetic diversity of dihaploids. (A) Circos plot showing gene density, variant density, diversity estimates (nucleotide diversity: *π* and Watterson’s theta: *θ_w_*), and population recombination rate (ρ). Positions on the DM reference genome are indicated in Mb and centromeres are marked in red. (A-1) Gene density per 1 Mb window. Variant density after (A-2) initial filtering (HF in Table 3) and (A-3) adjusted filtering parameters (HF+CF in Table 3), both within 1 Mb window. Nucleotide diversity in (A-4) HF and (A-5) HF+CF within 500 kb windows. Watterson’s theta in (A-6) HF and (A-7) HF+CF within 500 kb windows. (A-8) Historical recombination rate (*ρ*) of HF+CF set across 50-kb non-overlapping windows identified using FastEPRR 2.0. (B) Box and jitter plot showing *H_O_* across market classes. Each dot represents individual *H_O_* values, while asterisks denote the population’s expected heterozygosity. Letters above each box indicate groupings from post-hoc pairwise comparison using Dunn’s test. Market classes that share at least one letter are not significantly different (*p* > 0.05) and classes with no letters in common differ significantly (*p* < 0.05). (C) Information added by increasing sample size. We randomly subsampled our panel of dihaploids to examine the number of SNPs added when sample size increases. The X axis represents the size of the subsamples while the Y axis represents the total number of SNPs per subsample. Each point reflects the result of a single subsample. (D) Violin- and box-plot showing *π* (pink) and *θ_W_* (blue) for each market class. The central lines within each box indicate the median values, while the box edges represent the interquartile range (25^th^-75^th^ percentiles). Mean *π* values are marked by red circles while mean *θ_W_* values are indicated by red diamonds.

**Table 2.**
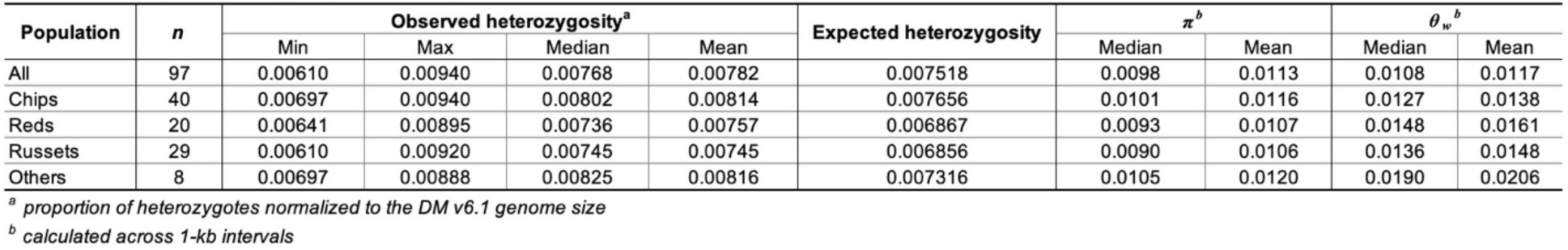
Summary of heterozygosity and diversity statistics across market classes.

**Table 3.** Summary of mean heterozygosity and diversity statistics across market classes under different filtering criteria.

| Population | n | Observed heterozygosity<sup>a</sup> | | | Expected heterozygosity | | | $\pi^b$ | | | $\theta_w^b$ | | |
| --- | --- | --- | --- | --- | --- | --- | --- | --- | --- | --- | --- | --- | --- |
|  |  | HF<sup>c</sup> | HF+CF<sup>d</sup> | hap52<sup>e</sup> | HF<sup>c</sup> | HF+CF<sup>d</sup> | hap52<sup>e</sup> | HF<sup>c</sup> | HF+CF<sup>d</sup> | hap52<sup>e</sup> | HF<sup>c</sup> | HF+CF<sup>d</sup> | hap52<sup>e</sup> |
| All | 97 | 0.00782 | 0.00088 | 0.00026 | 0.007518 | 0.000867 | 0.000254 | 0.0113 | 0.00299 | 0.00376 | 0.0117 | 0.00281 | 0.00339 |
| Chips | 40 | 0.00814 | 0.00092 | 0.00027 | 0.007656 | 0.000869 | 0.000253 | 0.0116 | 0.00306 | 0.00380 | 0.0138 | 0.00331 | 0.00400 |
| Reds | 20 | 0.00757 | 0.00085 | 0.00025 | 0.006867 | 0.000789 | 0.000233 | 0.0107 | 0.00293 | 0.00360 | 0.0161 | 0.00386 | 0.00466 |
| Russets | 29 | 0.00745 | 0.00085 | 0.00025 | 0.006856 | 0.000801 | 0.000237 | 0.0106 | 0.00289 | 0.00358 | 0.0148 | 0.00354 | 0.00428 |
| Others | 8 | 0.00816 | 0.00092 | 0.00027 | 0.007316 | 0.000836 | 0.000243 | 0.0120 | 0.00322 | 0.00390 | 0.0206 | 0.00494 | 0.00598 |

<sup>a</sup> mean proportion of heterozygotes normalized to the DM v6.1 genome size
<sup>b</sup> mean values calculated across 1-kb intervals
<sup>c</sup> properly paired reads, GATK hard-filtering parameters (HF), <10% missing rate prior BEAGLE v5.4 phasing and imputation (HF)
<sup>d</sup> primary alignment, GATKHF, MAPQ > 50, allele balance > 0.25, max depth = 2 x mean depth, and <1% missing rate prior phasing and imputation (HF+CF)
<sup>e</sup> loci in the HF+CF set that are 1-to-1 in at least 52 haplotypes (9,364 genes)

Expected heterozygosities (*H_E_*), calculated assuming Hardy-Weinberg Equilibrium for market classes and the population as a whole, were lower than mean and median *H_O_* (Table 2). However, the relative order of the market classes remained the same, with Chips exhibiting the highest *H_E_* and Russets exhibiting the lowest. Other measures of market class and population levels of diversity include, nucleotide diversity (*π*) and Watterson’s theta (*θ_w_*) (Table 2, and Figure 1A,D). Consistent with *H_O_* and *H_E_*, Chips exhibited the highest nucleotide diversity (mean *π* = 0.0116) and Russets showed the lowest (mean *π* = 0.0106). However, for *θ_w_*, Reds had the highest mean (*θ_w_* = 0.0161), while the Chips had a lower mean (*θ_w_* = 0.0138).

### Each additional dihaploid adds new variants

In order to guide our dihaploid extraction efforts, we investigated the information content each additional dihaploid sequence added to the data set by calculating the number of SNPs per subsample of the total data set (Figure 1C). Variation between SNP counts for different subsamples of the same size was as high as 12% for single clone samples in the Reds. However, the amount of variation decreased as sample size increased, with variation under 5% when six or more clones were included. The total number of SNPs in the subsamples increased as the number of dihaploids were added, and a limit was not reached for any of the three market classes. In general, while Russets and Reds had similar numbers of segregating sites, Chips had the most.

Although we attempted to maximize the number of tetraploid parents in our populations, we were limited by the inefficiency of dihaploid extraction (Busse et al., 2021). The P2DP included 1-5 dihaploids per parent. Dihaploids from the same tetraploid parent differed in 17- 24% of their SNPs, and limiting the data set to one individual per parent did not significantly change the summary statistic estimates.

### Haplotype diversity measures are impossibly high

We counted the number of alleles per locus, defined as a coding sequence (CDS) in the DM reference genome, using phased variants. Our initial analysis revealed a median of 16 alleles per locus (mean 26) (Figure 2A), with only 14 completely fixed CDSs, indicating regions of the genome with no variation among the 97 dihaploids. Conversely, for 44 CDSs we identified 194 alleles, meaning that every individual was heterozygous and no alleles were shared between individuals (Supplemental Table 7). Given that multiple dihaploids were derived from a single tetraploid mother, this outcome is impossible. To understand this further, we focused on dihaploids derived from the same tetraploid mother, targeting Caribou Russet, Red Norland, and W9968-5 from 3, 3, and 5 dihaploids per tetraploid parent, respectively, were sequenced (Figure 2). Each set of siblings exhibited some CDSs with complete heterozygosity and no shared alleles. Attempting to phase individual CDSs through manual curation, did not fix the problem.

**Figure 2.**
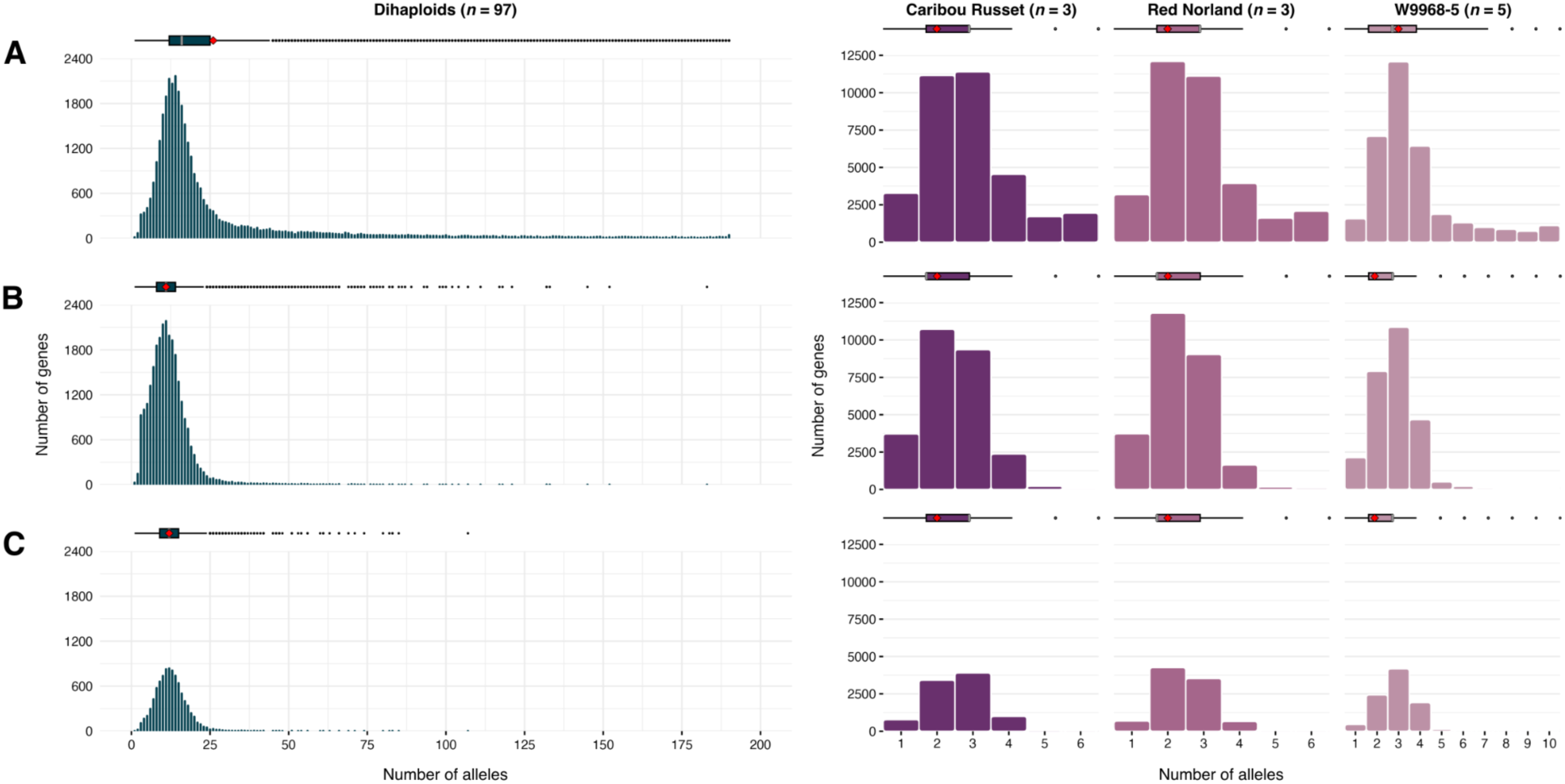
Allele distribution across dihaploids under different filtering criteria. Panels A-C show the allele distribution across all dihaploids (left-most column) and select dihaploids which share a single tetraploid parent (columns 2-4) across three different filtering criteria: (A) HF set: Properly-paired reads, MAPQ > 30, GATK hard filtering (HF), and a 10% missing rate prior to BEAGLE v5.4 phasing. The mean number of alleles per gene is 26 (red diamond) while the median is 16 (gray bar). (B) HF+CF set: More stringent filtering using primary alignments only, MAPQ > 50, HF, allele balance > 0.25, maximum depth ≤ 2x mean depth, excluding variants from low-complexity regions, and 1% missing rate before phasing. The mean and median number of alleles per gene is 11. (C) hap52 set: Same filtering as (B) but restricted to 9,364 genes from a synteny analysis that identified syntelogs that are 1-to-1 in at least 52 haplotypes. The mean and median number of alleles per gene in this set is 12. The 2^nd^ to 4^th^ columns display allele distributions for three dihaploids, each with at least three dihaploids sequenced that were generated from the same tetraploid mother. Each plot combines a histogram (bottom) of gene counts by allele number with a boxplot (top) summarizing the distribution of allele counts across genes (median: gray bar; mean: red diamond; outliers: circles). As filtering becomes more stringent from (A) to (C), the number of retained alleles per gene decreases.

To determine if sequencing errors caused this variability, we increased the stringency of variant filtering before phasing. We raised the mapping quality threshold from 30 to 50, used only primary alignments, limited read depth to twice the mean read depth, filtered out variant sites overlapping low-complexity regions, and decreased the missing rate from 10% to 1% (HF+CF set). These adjustments resulted in 3.5 million SNPs. Despite these changes, we still observed more than four alleles derived from a single tetraploid parent (Figure 2B) suggestive of structural variation within the tetraploid genomes, consistent with previous potato pan-genome analyses. To identify genes that are likely single copy in each haplotype within a tetraploid, we identified syntelogs present in individual haplotypes within DM, our 20 haplotype-phased dihaploids, and four tetraploid *de novo* haplotype-phased genomes (Figure 3). By restricting our analyses to 9,364 genes that are 1-to-1 syntenic in at least 52 haplotypes across DM, the dihaploids, and the tetraploids (hap52 set), the allele distribution became closer to the expected theoretical distribution, although some loci still exhibited more than four haplotypes (Figure 2C).

**Figure 3.**
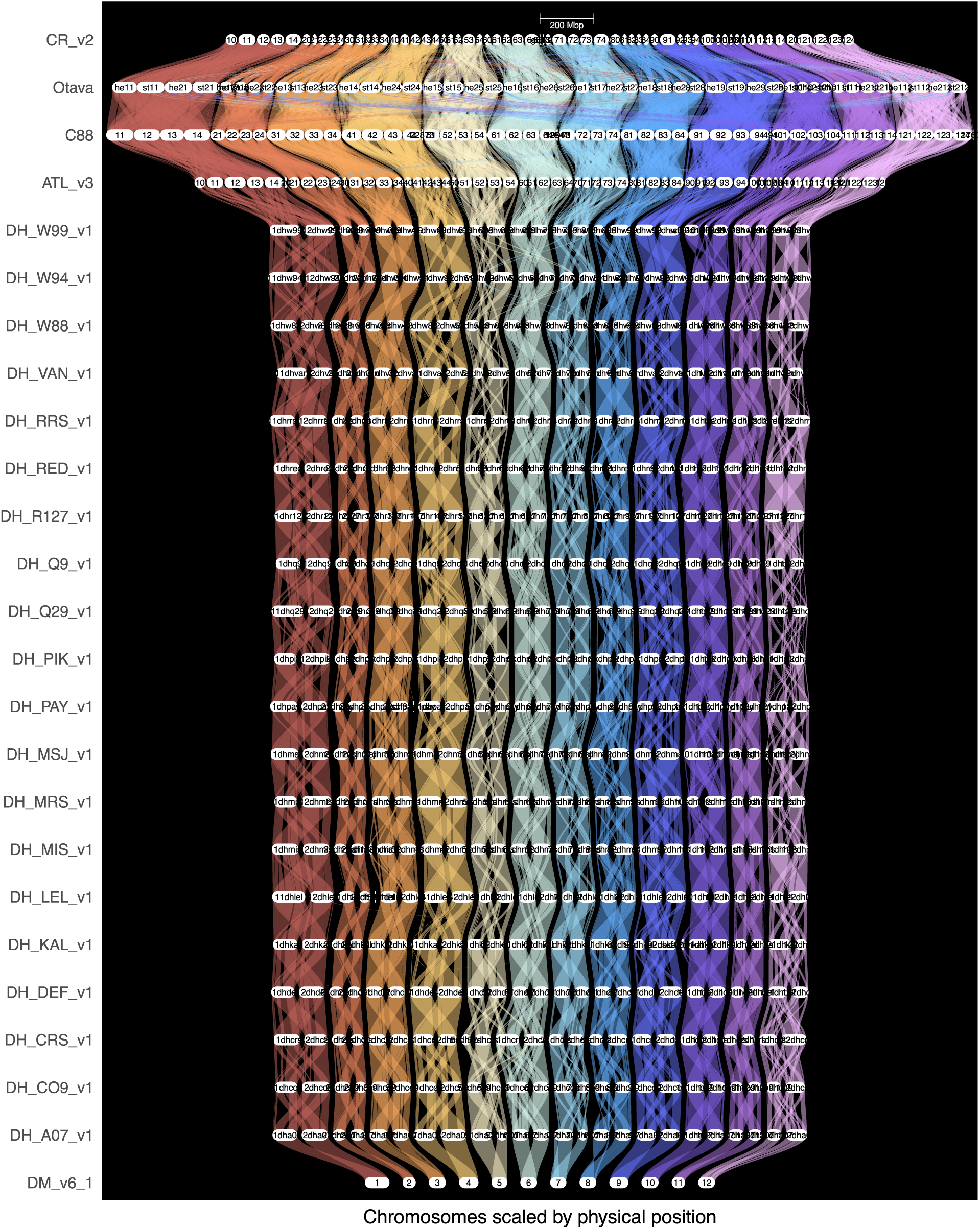
Synteny between genome assemblies of tetraploid potato, dihaploid potato, and the DM double monoploid using GENESPACE.

### Filtering criteria affect values of diversity statistics but not overall patterns

Overall, the diversity statistics calculated across market classes reveal similar patterns regardless of the filtering criteria applied (Table 3). Observed heterozygosity (*H_O_*) shows a consistent trend across market classes, with the highest value for Chips and slightly lower values for Reds and Russets. Additionally, *H_O_* consistently exceeds expected heterozygosity across all market classes, regardless of the filtering criteria used. For nucleotide diversity (*π*), the HF+CF set shows lower values across all market classes when compared to the HF set. In contrast, the hap52 set yields slightly higher *π* values, ranging from 0.0036 in Russets to 0.0038 in Chips.

Similarly, Watterson’s theta (*θ_w_*) follows the same pattern with a slightly higher diversity estimate in the hap52 set compared to HF+CF. The *θ_w_* values in the HF+CF set range from 0.00331 in Chips to 0.00386 in Reds, whereas the hap52 filter ranges from 0.004 in Chips to 0.00466 in Reds.

### Role of CNVs in haplotype diversity

Given the presence of paralogs and observed genetic variability, copy number variations (CNVs) likely account for some of the alleles observed among dihaploids. An initial read-depth-based CNV analysis revealed that 14.7% to 19.4% of genes in individual dihaploids were either duplicated or deleted, with an average of 16.5% (Supplemental Table 8). Specifically, 31% of genes exhibited deletions, including 18.7% with homozygous deletions, while 7% were duplications, and 7.3% showed both duplications and deletions. For dihaploids derived from the W9968-5 tetraploid parent, a chi-square test on read-depth based CNVs demonstrated a strong association between the number of alleles (categorized as 1-4 and >5) and the presence of CNVs (p = 2.2 × 10⁻^16^; Supplemental Table 9).

To conduct a more comprehensive analysis of CNVs across the genome, duplications and deletions were identified using three structural variation (SV) callers: Manta, DELLY2, and Smoove. CNVs were merged using SURVIVOR if at least two variant callers supported the CNV, resulting in a non-redundant set of 43,477 deleted and 8,814 duplicated regions (Figure 4). From the 40,383 genes analyzed in the initial read-depth-based CNV analysis, 22,376 (55.4%) overlapped with the CNVs identified by the SV callers. The analysis revealed varying levels of CNV sharing across the population. CNVs shared by at most 10 individuals included 1,812 duplications and 11,637 deletions, while CNVs shared by at least 91 individuals included 503 duplications and 3,936 deletions.

**Figure 4.**
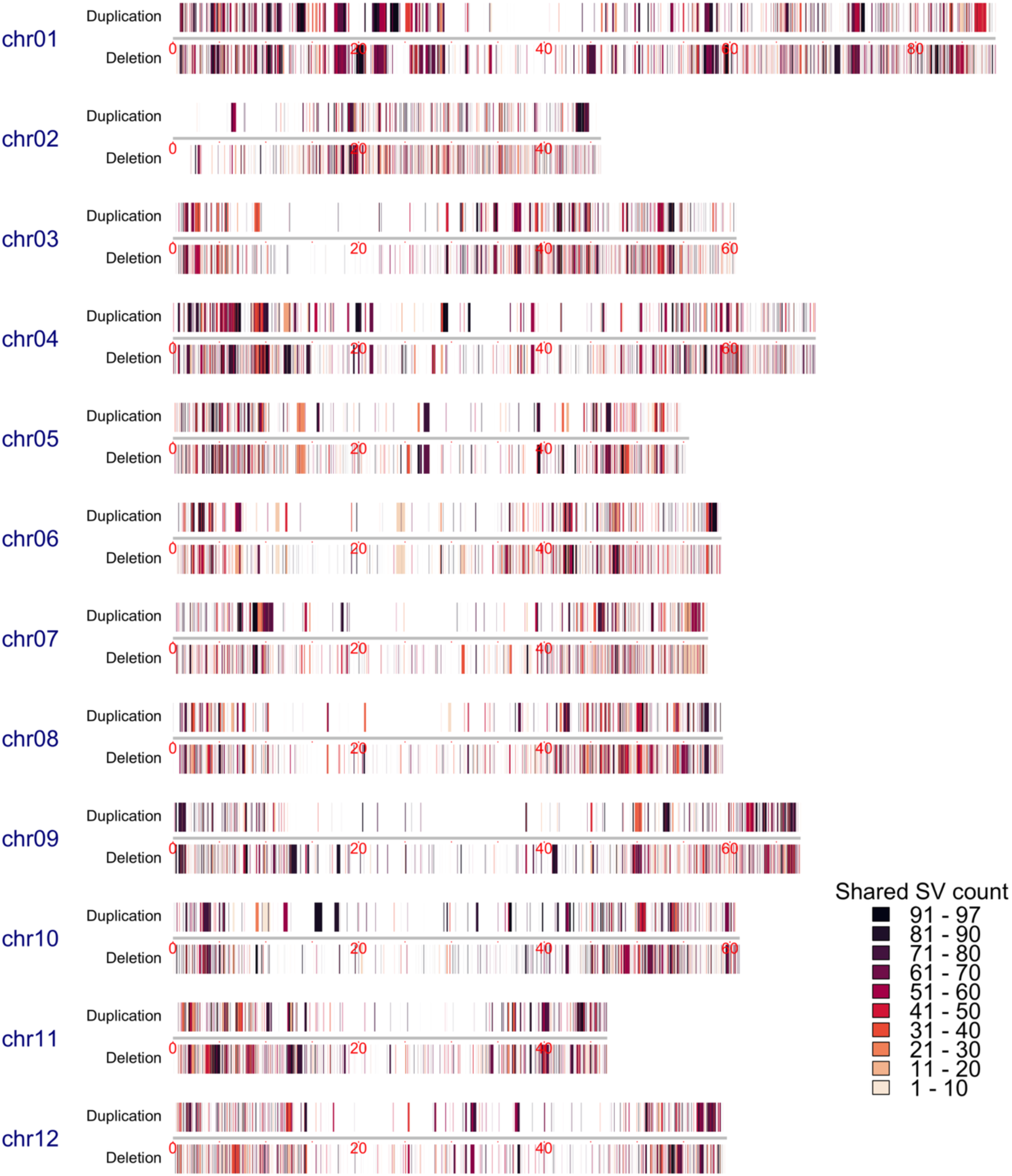
Distribution of shared duplications and deletions among the dihaploids. Chromosome distances are marked in Mb. Duplications are depicted above the chromosome, with color indicating the number of individuals they are found in, while deletions are similarly represented below the chromosome.

### Dihaploids have long linkage blocks

Potato exhibits low genome-wide recombination (*ρ = 2 kN_e_ r*) compared to other plants (Figure 5; 1001 Genomes Consortium, 2016; Zhang et al., 2021; Guo et al., 2019; Yang et al., 2021; Sun et al., 2020; Wang et al., 2018; Wu et al., 2020; Liu et al., 2022; Lozano et al., 2021; Cai et al., 2021; Zhou et al., 2020). The historical recombination rate, reflecting a history of recombination between unique haplotypes, of the sequenced dihaploids ranged from 0 to 339.58 per kb with a genome-wide average of 1.42 per kb (Figure 5, Supplemental Figure 4 and Supplemental Table 10). When analyzed by market class, Chips had a higher genome-wide average *ρ* of 1 per kb, while Reds and Russets had lower averages at 0.65 per kb. Chromosome 4 displayed the highest *ρ* across all market classes except in Russets with an average of 2.05 per kb in the combined population, and 1.55 per kb, 0.90 per kb, and 0.87 per kb in Chips, Reds, and Russets, respectively. In contrast, the recombination rate was lowest on chromosome 2 in the combined population, with the lowest rates observed in Russets (*ρ* = 0.43 per kb). A weak negative correlation was observed between *π* and *ρ* (Spearman’s ρ = -0.352; *p* < 0.0001) across genomic intervals in all sequenced dihaploids.

**Figure 5.**
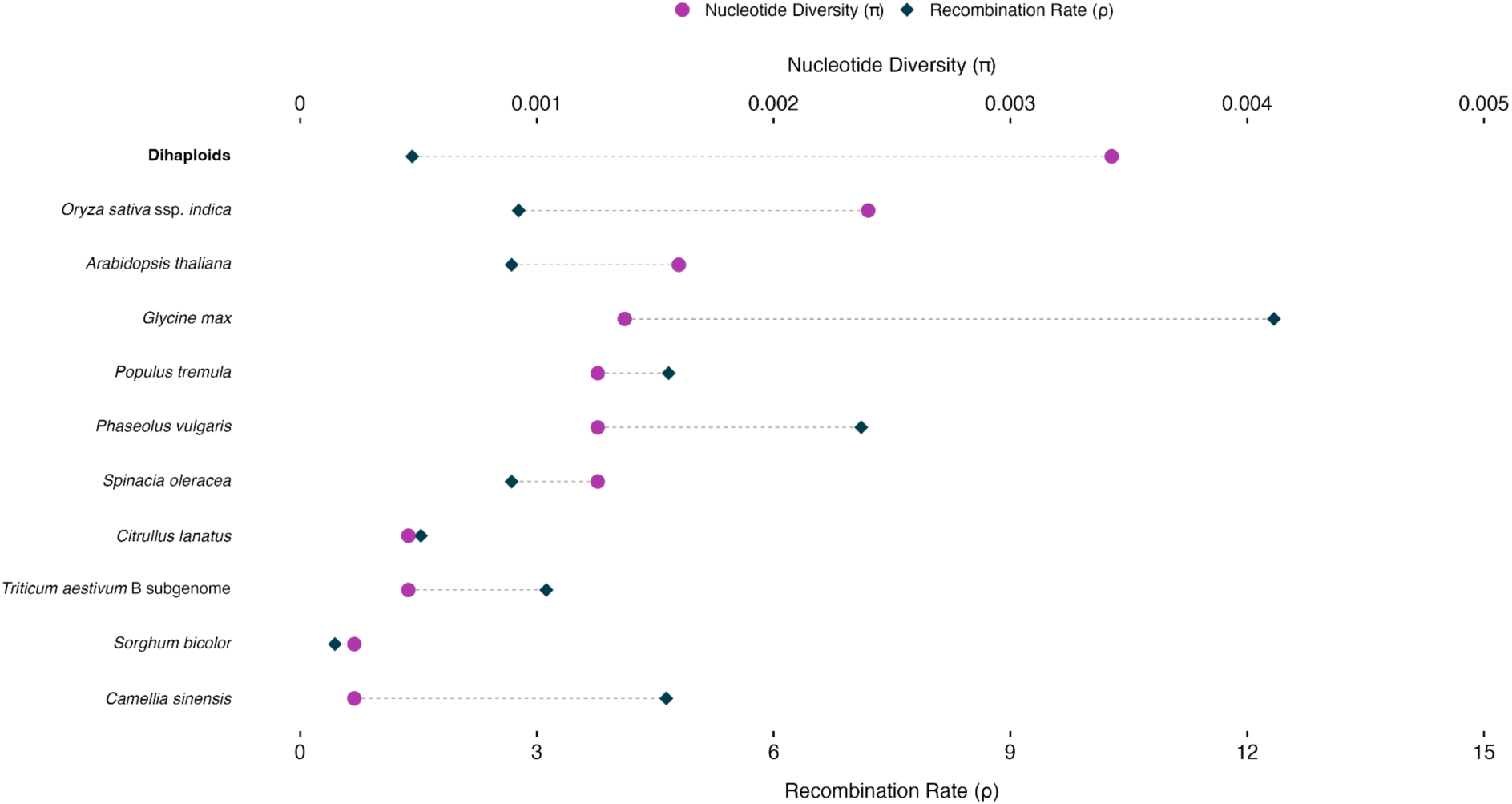
Recombination rate (*ρ*/kb, lower axis, blue diamond) and nucleotide diversity (*π,* upper axis, purple circles) of potato dihaploids compared to other plants. (Brazier & Glémin, 2024; 1001 Genomes Consortium, 2016; Zhang et al., 2021; Guo et al., 2019; Yang et al., 2021; Sun et al., 2020; Wang et al., 2018; Wu et al., 2020; Liu et al., 2022; Lozano et al., 2021; Cai et al., 2021; Zhou et al., 2020)

### Allelic diversity at StCDF1, the potato maturity gene

A total of 15 *StCDF1* alleles were predicted among the 194 dihaploid haplotypes based on 118 phased SNPs in the HF marker set and indel variants in the second exon. The most common allele, found in 62 haplotypes (red labels in Figure 6), was *StCDF1.3*, which conveys early maturity under long days due to an 865 bp transposon insertion that disrupts the FKF1 protein binding domain and the long non-coding RNA *StFLORE* (Kloosterman et al. 2013; Ramírez Gonzales et al. 2021). Alleles *StCDF1.2T* and *2C*, shown in blue and purple, respectively, cluster with *StCDF1.3* because all three have identical phased SNP haplotypes (equal to the DM1-3 allele). The *StCDF1.2* alleles have a 7 bp insertion relative to the DM reference, hypothesized as the remnant of the transposon excision from *StCDF1.3* (Kloosterman et al. 2013). A [T/C] SNP exists between the *2T* and *2C* alleles in the 7 bp insertion. One allele, from the European variety ‘Malou’, contained the full transposon but differed from *StCDF1.3* at three SNPs. More common than *StCDF1.2* was *StCDF1.4* (Gutaker et al. 2019; green in Figure 6), which contains a different 7 bp insertion in a different genetic background. Both *StCDF1.2* and *StCDF1.4* convey early maturity due to disruption of the FKF1 binding domain, but the effect is not as strong as *StCDF1.3* (Hoopes et al. 2022; Caraza-Harter and Endelman 2022; Ma et al. 2025). The five largest clusters with black labels in Figure 6 contain at least one haplotype from the assemblies, and the predicted CDF1 proteins contain the FKF1 binding domain (i.e., type *StCDF1.1*), which promotes late maturity under long days.

**Figure 6.**
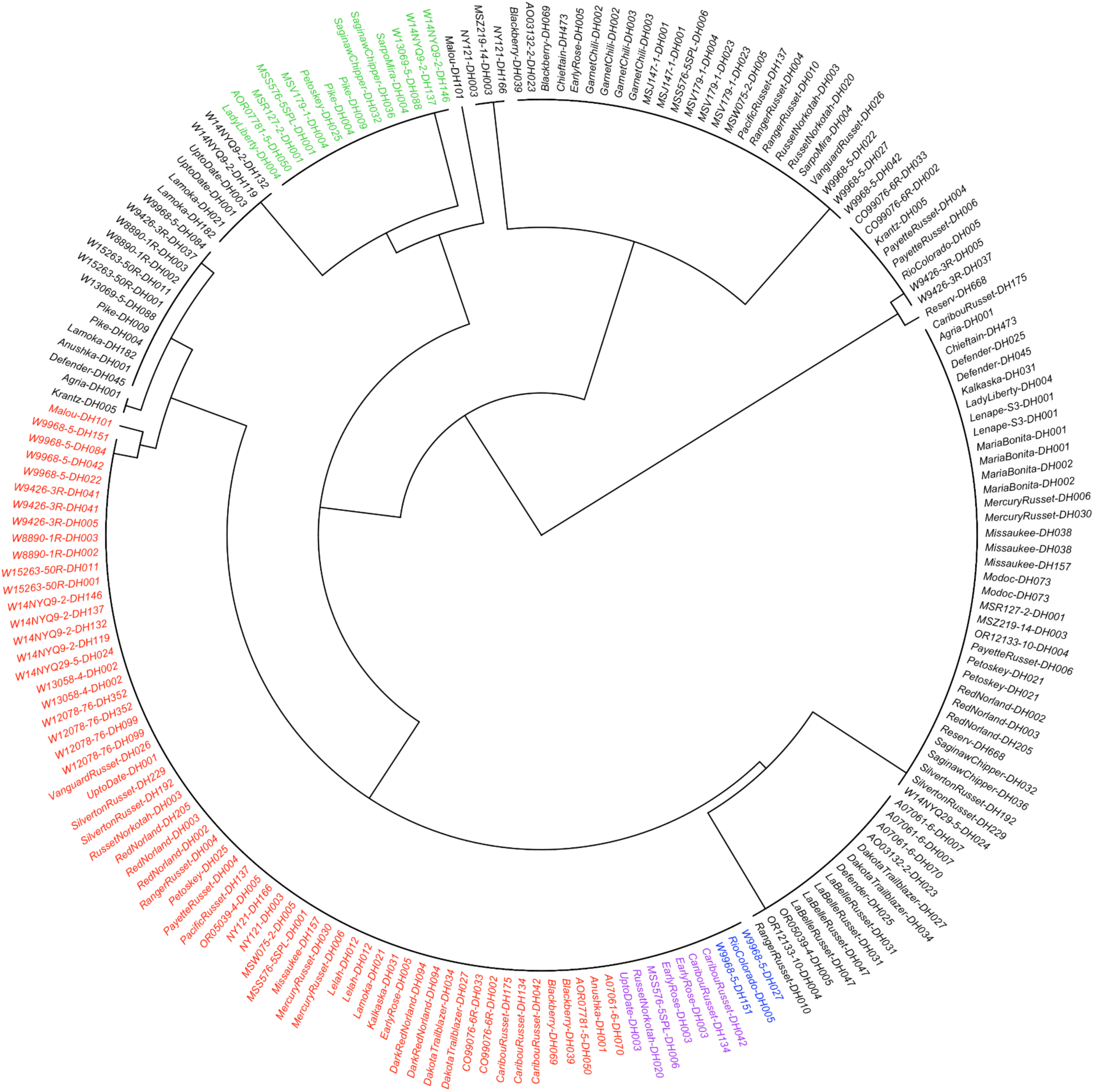
UPGMA clustering of phased SNP haplotypes (HF marker set) in the 97 dihaploids. Red labels are *StCDF1.3*; blue are *StCDF1.*2*T*; purple are *StCDF1.2C*; green are *StCDF1.4*; and black are *StCDF1.1*.

## Discussion

### A foundation for diploid potato breeding

Development of a dihaploid population is a critical step toward establishing a diploid breeding framework within the U.S. context (Jansky et al., 2016). We developed three sets of dihaploids, which are available through the U.S. potato genebank and have already been distributed to several public and private sector breeding programs. The first is derived from round white potatoes with high specific gravity used for making potato chips. The second is derived from oblong russeted potatoes with high specific gravity used for French fries and other processed potato products. The third is derived from red skinned white fleshed lower density potatoes consumed as fresh vegetables. The clones described here contain alleles for crucial potato traits as shown through visible phenotypes observed in the dihaploids such as russeting, shape (oblong and round), maturity, and skin color. They also, presumably, contain extensive deleterious alleles. Most of the 97 dihaploids produce tubers and are female fertile, but only one is male fertile and most exhibit low yield and reduced vigor. Previous research in diploid potatoes suggests that the potato germplasm is rife with deleterious and dysfunctional alleles (Zhang et al., 2019; Zhang et al., 2021; Wu et al., 2023). We expect an even higher frequency of deleterious alleles in the autopolyploid parents of the 97 dihaploids due to relaxed selection (Monnahan & Brandvain, 2020; Booker & Schrider, 2023). As demonstrated by Zhang et al. (2021), a design breeding approach based on sequence information could facilitate the establishment of diploid potato varieties on a rapid time scale. If this dihaploid population is used as a starting point for population development, sparse genotyping and imputation (Swarts et al., 2014; Bradbury et al., 2022) will allow for haplotype reconstruction in breeding populations.

### Paralogs prevent a single unified interpretation of the sequence data

Generation of whole genome assemblies from dihaploids with only two haplotypes reduces the level of complexity and cost relative to a genome assembly of a tetraploid. For 20 dihaploids, we generated phased, haplotype-resolved assemblies with high contiguity as shown by assembly metrics and representation of BUSCO orthologs. Annotation of the dihaploids using an identical workflow that included full-length cDNA sequences from leaves and tubers of each dihaploid enabled identification of synthetic genes within and between haplotypes of the dihaploids, cultivated tetraploids, and the reference DM genome. From synteny analysis of 57 potato haplotypes using DM as the reference, we identified 1,154 syntelogs present in all 57 haplotypes, 28,468 syntelogs present in 2 to 56 haplotypes, and 3,279 DM genes not found in syntenic groups. This structural variation at the synteny level is consistent with previous findings in cultivated tetraploid potato (Hoopes et al. 2022) and due to deletions, duplications, and introgressions. However, a subset of structural variation may be attributable to assembly errors or annotation artifacts.

The extensive structural variation (Pham et al., 2017; Bozan et al., 2023) in the dihaploids presents a challenge for interpretation of sequencing data as paralogs introduce additional alleles that can mimic haplotypes and inflate diversity metrics (Bohutínská et al., 2023). Despite stringent variant filtering, we observed more than four alleles per locus in dihaploids derived from single tetraploid mothers. Only filtering out all heterozygous loci prevented “excess” alleles in dihaploids derived from a single tetraploid parent, however, we know the dihaploid lines are not isogenic from genotyping. We therefore conclude that these additional haplotypes are not artifacts of sequencing, but rather reflect real genomic complexity contributed by paralogs.

The presence of paralogs can inflate perceived genetic diversity as they are conflated with true allelic variation. To eliminate the paralogs from the analysis, we generated a data set consisting only of genes which were present in single copy in at least 52 of 57 haplotypes identified through synteny analyses of the DM reference genome, 20 phased dihaploid genomes, and four tetraploid haplotype-phased genome assemblies [Castle Russet (Hoopes et al., 2022), Otava (Sun et al., 2022), Cooperation-88 (Wang et al., 2022), Atlantic (Hoopes et al., 2022)].

These 9,364 single-copy genes are presumably under the greatest evolutionary constraint and therefore the least variable across these 57 haplotypes. While the original analysis is likely an overestimation of diversity due to paralogs, the analysis based on the 1-to-1 syntelog set is likely excessively conservative. Therefore, we have chosen to present both analyses. It is worth noting that even the conservative analysis results in higher nucleotide diversity than that of wheat, sorghum, cultivated and landrace watermelon, cotton, and fresh-market tomato (Zhou et al., 2020, Boatwright et al., 2022, Guo et al., 2019, Li et al., 2023, Bhandari et al., 2023) and most patterns of diversity within the dihaploids remain constant regardless of how the data were analyzed.

### Elucidating the tetraploid potato genome

In addition to forming a base population for diploid breeding, these dihaploids provide a powerful tool for genetics. In this study, we used sequence analysis technologies designed for diploids to reveal haplotypes within cultivated potatoes, which provide insight into tetraploid genomes and evolution. We found high levels of genetic diversity in the population (*π, θ_w_,* expected heterozygosity). However, observed heterozygosity, H_O_, was even greater (mean Tajima’s D for HF+CF set = 0.2). In the U.S. there is a history of intentional breeding for heterozygosity in potatoes (Hermundstad & Peloquin, 1987), but this pattern extends beyond the U.S. as observed heterozygosity in excess of expected heterozygosity has been reported across *S. tuberosum* Group *Chilotatum*, unlike *S. tuberosum* Group *Andigenum* where there is evidence of mild inbreeding (Tuttle et al., 2024). This is likely due to selection against homozygosity in the face of inbreeding depression, which has been repeatedly demonstrated in potatoes (Zhang et al., 2019; Zhang et al., 2021; Manrique-Carpintero et al., 2018; Achakkagari et al., 2022; Hoopes et al., 2022). The combination of high levels of population diversity and intentional or incidental balancing selection by breeders throughout time likely both contribute to the excess of observed heterozygosity in U.S. cultivated potatoes.

The high nucleotide diversity observed in our study reflects the history of the tetraploid mothers. Autotetraploids present a larger mutational target (Meirmans et al., 2018), with more copies of each chromosome, increasing the chances of mutation accumulation. Furthermore, in potato, introgression from numerous wild relatives has contributed substantially to genetic diversity in potato (Hoopes et al., 2022; Meng et al., 2022; Hardigan et al., 2017, Vos et al., 2015, Anglin et al., 2024; Sun et al., 2025). Much of this diversity is maintained in the population because the effects of selection are reduced in autopolyploids, resulting in the retention of deleterious alleles (Monnahan and Brandvain, 2020). Inefficient selection in tetraploids may also contribute to the prevalence of paralogs and other CNVs in the dihaploids.

We used multiple measures for population level diversity. Watterson’s theta (*θ_w_*) reflects the frequency of segregating sites. These segregating sites arose at one of three points in potato’s evolutionary history: before speciation, as mutations in *S. tuberosum*, or in wild relatives followed by introduction to cultivated potato through introgression. Nucleotide diversity (*π*) reflects the distribution of these variants in the population. An excess of low frequency variants, of the type associated with bottlenecks, produces high values of *θ_w_* but low values of *π*. We observe high values of both *θ_w_* and *π,* suggesting extensive introduction of diversity through mutation and introgression but also distribution of that variation across the gene pool. This indicates that the variation is not particularly recent and has had time to spread in the population, a conclusion consistent with the observations of shared introgressions in the potato pangenome (Meng et al., 2022; Hoopes et al., 2022).

Another method for describing diversity is historical recombination rate (*ρ = 2kN_e_r*), which reflects a history of recombination between unique haplotypes (Hill, 1981; Pritchard & Rosenberg, 1999; Slatkin, 2008; Hayes et al., 2003). For a high diversity species like potato we would expect a high *ρ*. However, *ρ* in potato is low. Only sorghum and watermelon exhibit comparatively low *ρ*, but they have correspondingly low nucleotide diversity (Figure 5). This suggests that while diversity is high in potato, much of that diversity is in linkage blocks resulting in low haplotype diversity (Sun et al., 2025).

The *ρ* metric depends on both effective population size (*N_e_*) and the average recombination frequency (*r*) for the window. The latter has been estimated in many genetic mapping studies (Massa et al., 2015; Felcher et al., 2012; da Silva et al. 2017; Clot et al., 2024; Manrique Carpintero et al., 2016; Manrique Carpintero et al., 2015; Prascher et al., 2014; Park et al., 2021). Most estimates fall in 1.1-1.5 cM/Mb, although extreme values of 0.9 (Manrique Carpintero et al., 2016; Park et al., 2021) and 3.07 cM/Mb (Bourke et al., 2015) have been reported. From a survey of other species (e.g., rice, soybean, sorghum), recombination frequency in potato is typically 2–3 times lower (Supplemental Table 11). A reduced recombination rate is an essential component in genome stabilization immediately following whole-genome duplication events in *Brassica* (Gonzalo et al., 2023; Morgan et al., 2021). Such a reduction allows polyploids to manage chromosomal pairing complexity (Otto et al., 2007). If reduced recombination followed the potato whole genome duplication events, this could partially explain the gap between *ρ* and *π*.

An alternative explanation for low measures of *ρ* is a small effective population size. However, *θ_w_*, for which we observe high values, is also dependent on effective population size (*θ= 2kN_e_µ*). Some of the discrepancy between *ρ* and *θ_w_* could be explained by departure from neutral expectations. A bottleneck and migration have been proposed as explanations (Sun et al., 2025), and as all of these clones are derived from breeding programs, selection and assortative mating have also certainly taken place. The equation for *θ_w_* suggests a high mutation rate as another possible explanation for the difference in these two measures of diversity. Potatoes can reproduce clonally resulting in long generation times and heritable somatic mutations. The smaller number of generations provides few opportunities for recombination, but the long generation time provides extensive opportunity for mutation accumulation. Although we do not have a good estimate for the somatic mutation rate in potato, there is evidence of extensive somatic mutation in tissue culture (Bozan et al., 2023) and the field (Waterer et al., 2011; Miller et al., 2004; Nassar et al., 2011). This combination likely contributes to the observed discrepancy.

Much like limited generations, introgression can result in extended haplotype blocks, because an introgressed haplotype has had fewer opportunities to undergo recombination in the recipient genome than syntenic regions without introgression (Vernot & Akey, 2014). The history of potato is characterized by extensive introgression from over 20 species (Hoopes et al., 2022; Meng et al., 2022; Hardigan et al., 2017, Vos et al., 2015, Anglin et al., 2024; Sun et al., 2025); this may create a large enough effect to affect *ρ*.

Recombination rate, mutation rate, generation time, and introgression all likely play some role in the discrepancy between *ρ* and *θ_w_* . Further work is needed to determine the relative contribution of each factor.

### Next steps for diploid potato breeding

We have developed a population of 97 dihaploid potatoes which contain many of the alleles from U.S. commercial tetraploid potatoes. It is unlikely that these dihaploids contain the complete set of U.S. commercial haplotypes as each sequence added to our panel generated new previously unrecorded variants, suggesting that additional dihaploids would likely further expand our allele set. However, the majority of genes have a median of 16 and a mean of 26 haplotypes in our panel (Figure 2A). This distribution is right skewed, suggesting that for much of the genome most alleles have been found; however, highly variable regions likely contain alleles that have not been sampled. Observations of the phenotypes in the set of dihaploids and breeding efforts thus far (Alsahlany et al. 2021, Marand et al. 2019, Song & Endelman, 2023), suggest that we have captured the majority of the crucial alleles to the U.S. potato breeding and processing industry. However, alleles outside this collection will likely be necessary, and require introgression from related germplasm. At minimum, diploid male fertility must be brought in from another background.

The primary challenge to breeders establishing inbred lines will be recessive and partially recessive deleterious alleles, which cause inbreeding depression (Alpers & Jansky, 2019; de Vries et al., 2023). Large effect alleles may be purged through design breeding (Zhang et al., 2021), but purging must be carefully managed to avoid compromising diversity, which is essential for adaptability and resilience (Morrell et al., 2012). Furthermore, purging all or even most deleterious alleles on a reasonable time scale is impossible (Labroo et al., 2023). While complementation, through hybrid breeding is the long-term goal (Lindhout et al., 2011; Jansky et al., 2016; Zhang et al., 2021; Bradshaw, 2022; de Vries et al., 2023), preliminary population improvement to break up long linkage blocks with high deleterious load may be necessary.

Breeding for maturity, via selection of specific *StCDF1* alleles, touches on many of the aforementioned themes. Our results show there are two common allelic groups in North American varieties: full-length *StCDF1.1* alleles and the truncated allele *StCDF1.3.* Balancing selection is the best explanation for this result, as clones homozygous for *StCDF1.1* are generally too late-maturing (under long days), while clones homozygous for *StCDF1.3* are too early (Caraza-Harter and Endelman 2022; Ma et al. 2025). The *StCDF1* genotype 1/3 produces a desirable maturity, but this combination is not efficiently maintained under sexual reproduction in a single breeding pool, as 50% of all offspring from 1/3 x 1/3 crosses have undesirable genotypes. However, in the context of diploid hybrid breeding, fixing one pool for *StCDF1.1* and the other for *StCDF1.3* leads to the desired hybrid genotype. Another direction may be to fix the intermediate-maturity alleles *StCDF1.2* or *StCDF1.4*, which were more rare in the 97 dihaploids (Figure 6), but overly rapid fixation could lead to linkage drag from deleterious alleles at nearby loci.

Consolidating the allelic diversity of tetraploid varieties into a diploid form allows for a more targeted exploration of complex traits. By reducing some of the challenges associated with autopolyploidy, dihaploids provide a streamlined approach to identifying and fixing desirable traits. This population provides a starting point for breeders interested in developing new diploid potato varieties, especially those addressing U.S. markets. Furthermore, the genomic resources we generated for this population allow for the implementation of design breeding, facilitating rapid genetic gain. This combination positions us well to reinvent potato as a dynamic crop able to quickly respond to coming changes in the environment and consumer demand.

## Materials and Methods

### Plant materials and whole genome resequencing

We pollinated 58 tetraploid potato cultivars and advanced breeding lines with the *Solanum phureja* line, IVP101, to make dihaploids (Busse et al., 2021; Hougas and Peloquin 1957; Hougas and Peloquin 1958; Hougas et al. 1958). Putative polyploids were removed based on seed spots (Uijtewaal, Huigen and Hermsen 1987), plant vigor, flow cytometry, chloroplast counting (Kramer and Bamburg 2019), and genotyping (see https://github.com/jendelman/polyBreedR, Alsahlany et al., 2019). Dihaploids were assessed in the greenhouse at the University of Minnesota, University of Wisconsin, Michigan State University, University of Maine, Oregon State University, and the Pepsi Potato Research Facility in Rhinelander Wisconsin. Only those that produced tubers, flowers, and true potato seed upon pollination were considered for sequencing. Dihaploids were further prioritized for sequencing based on pedigree and other known relationships (Bali et al., 2018; Hirsch et al., 2013).

We collected leaf samples from dihaploids grown in tissue culture for DNA extraction. The DNA was extracted using the Qiagen DNeasy Plant Mini Kit (Hilden, Germany) according to the manufacturer’s protocol. After extraction, we resuspended the genomic DNA in distilled, deionized water and shipped it on dry ice to the University of Minnesota Genomics Center for library preparation and sequencing. Libraries were constructed using the Illumina TruSeq Nano DNA Library Preparation Kit and then sequenced to a target depth of 20x coverage on an Illumina NovaSeq 6000 platform in paired end mode to 150 nt (San Diego, CA).

### Quality control of raw sequence data

The quality of the raw sequence data was evaluated using FastQC v0.11.9 (Andrews, Simon, 2010) to check for adapter contamination, %GC content, and overrepresented sequences. We then reviewed aggregated results using MultiQC v1.21 (Ewels et al., 2016) to streamline the evaluation of multiple FastQC reports. Low-quality reads and adapter sequences were then trimmed using Trimmomatic v0.39 (Bolger et al., 2014 https://github.com/usadellab/Trimmomatic). Illumina adapter sequences, included with Trimmomatic, were used to identify adapter contamination. The adapter trimming process was carried out in palindrome mode, allowing two seed mismatches, with a palindrome clip threshold set at 30, a simple clip threshold of 10, and a minimum adapter length of seven nucleotides required for clipping. Additionally, we trimmed paired-end reads for low quality scores (Phred score < 3) in the leading and trailing sequences, and within a sliding window of size four nt.

Reads were removed if their minimum length fell below 36 bp. Only paired sequences, where both forward and reverse reads remained post-trimming, were subsequently mapped to the reference genome.

#### Processing of filtered reads and ploidy estimation

Paired sequences were aligned to the double monoploid *S.tuberosum* Group Phureja DM 1-3 516 R44 v6.1 reference genome (Pham et al., 2020) using the BWA-MEM algorithm in the Burrows-Wheeler Aligner v0.7.17-r1198-dirty (Li & Durbin, 2010), set to default settings. After alignment, the resulting binary alignment (BAM) files were sorted by coordinates and indexed using SAMtools v1.17 (Danecek et al., 2021). We added read groups into the sorted BAM files using AddOrReplaceReadGroups command in Picard tools v3.0.0 (Broad Institute, 2019 https://broadinstitute.github.io/picard/), included with GATK v4.5.0.0 (Van der Auwera et al., 2020). PCR duplicates were then identified and marked using the MarkDuplicates command in Picard tools. We estimated ploidy levels from the sorted and deduplicated BAM files, filtering for reads that were properly paired and had a minimum quality score of 30, using SAMtools (v1.17; Danecek et al., 2021). The filtered BAM files were then used to determine ploidy using nQuire (Weiß et al., 2018; https://github.com/clwgg/nQuire), setting the minimum coverage at 20 per position.

#### Variant discovery

Single nucleotide polymorphism (SNP) discovery in each dihaploid was performed using the HaplotypeCaller tool in GATK (v4.5.0.0; Poplin et al., 2018), setting the minimum mapping quality to 50 and the minimum base quality score to 30. We configured the variant calling for diploid samples in emit-ref-confidence genomic variant call format (GVCF) mode and restricted to primary alignments by applying the PrimaryLineReadFilter flag. The GVCF generated for each sample was combined per chromosome using the CombineGVCFs tool, followed by joint genotyping of all samples per chromosome using GenotypeGVCFs. We then restricted the dataset to biallelic SNPs, excluding all multiallelic sites. Variant sites were further filtered following GATK’s best practices on hard-filtering, which include a quality by depth greater than 2, quality score greater than 30, strand odds ratio less than 3, Fisher strand less than 60, mapping quality greater than 40, mapping quality rank sum test greater than -12.5, and ReadPosRankSum greater than -8. We applied additional genotype-level filtering criteria, where genotypes were assigned as missing if the quality score was below 30 and allele depth was less than 10. Sites with a missing rate of less than 10% were retained for further analysis. The filtered VCF file was then phased using BEAGLE v5.4 (Browning et al., 2021).

Structural variants were called for each dihaploid using the duplicate marked, coordinate-sorted BAM files. These were then genotyped at the population-level using Delly v1.2.6 (Rausch et al., 2012) and Lumpy via Smoove v0.2.8 (Layer et al., 2014; Pederson et al., 2020 https://github.com/brentp/smoove/). Only variants that passed the filtering criteria of each SV caller were used in subsequent analysis. To create dihaploid-specific SVs, calls from each tool were merged on a per-individual basis using SURVIVOR v1.0.7 (Jeffares et al., 2017 https://github.com/fritzsedlazeck/SURVIVOR), requiring breakpoints within 1 kb, at least two of the three callers supporting the variant, and consistent SV type and strand. A non-redundant variant call set was then created by merging SVs across the dihaploids using BCFtools v1.17 (Danecek et al., 2021). Additionally, we used Duphold v0.2.3 (Pedersen & Quinlan, 2019 https://github.com/brentp/duphold) to obtain read depth for regions of overlapping SVs .

#### Genome-wide patterns of heterozygosity, nucleotide diversity, Watterson’s theta, and recombination rate

Observed heterozygosity was calculated in Python as the proportion of heterozygous loci across samples. For each individual, we counted heterozygous calls from the VCF file and divided by the genome size of DM v6.1 to normalize values. Expected heterozygosity (H_E_), under Hardy-Weinberg equilibrium, was calculated as one minus the sum of the squared allele frequencies for each locus. We then average H_E_ values across all loci to obtain a genome wide estimate.

To examine heterozygosity differences across subpopulations, we used Kruskal-Wallis Rank Sum test, followed by post-hoc pairwise Dunn’s test with Benjamini-Hochberg correction using the *kruskal.test* and *dunn.test* functions in R (v4.3.0). Non-parametric methods were selected based on non-normal distribution and homogeneity of heterozygosity data, confirmed by the Shapiro-Wilk normality test *shapiro.test* function in R (*p*-value: Chips = 0.000271, Reds = 0.002460, Russets = 0.000144, Others = 0.000468) and equal variances verified at a 95% confidence level using Levene’s Test *leveneTest* function in R (*p*-value = 0.08966).

Watterson’s theta was calculated using SNP density data generated with VCFtools at 1- kb intervals (v 0.1.17; Danecek et al., 2011). For each market class, we calculated the harmonic number *h* as the sum of the reciprocals from 1 to *n* - 1, where *n* is twice the sample size.

Watterson’s theta for each interval was then calculated by first dividing the number of segregating sites by the harmonic number and then by interval size. Population-level *θ_w_* was obtained by averaging interval estimates across the genome.

Nucleotide diversity was calculated for each market class using VCFtools (v 0.1.17; Dancek et al., 2011) at 1-kb and 50-kb intervals. Historical recombination rates were then estimated using FastEPRR 2.0 (Gao et al., 2016), with non-overlapping windows of 1-kb and 50- kb. To align the recombination rate and nucleotide diversity estimates across the same intervals, we set *erStart = 1* to enable direct comparison between the two metrics.

#### Allele analysis

We developed a custom Python script to assess the additional information added with each new sequence. For each market class, we sampled one individual at random and recorded the number of SNPs. Then a second individual was added at random and new SNPs were counted. This was continued until all sequences were added. This was repeated 40 times for each market class and the number of SNPs added with each individual was plotted against the population size.

The number of alleles per CDS was counted using a custom Python script designed to process phased variants from locus-specific VCF files. This script determined the number of unique alleles for each CDS across all dihaploids and within each market class. Only identical alleles were counted as the same.

#### StCDF1

Pairwise Hamming distances were calculated for the 194 haplotypes of the 97 dihaploids using the 118 SNPs in Soltu.DM.05G005140 with the HF filtering parameters. The dendrogram was constructed using R/hclust for hierarchical clustering with method “average”. Haplotypes were classified as 1, 2T, 2C, 3, or 4 based on alignment (BWA-MEM) of the Illumina WGS data to reference sequences for *StCDF1.1* (scaffold1389) and *StCDF1.3* (scaffold1390) from the Atlantic reference genome (Hoopes et al. 2022). Variant discovery and genotype calling were done with freebayes (Garrison and Marth 2012). Alleles 2T, 2C, and 4 were determined based on the sequence of the 7 bp insertion in the *StCDF1.1* alignment.

### Dihaploid Genome Sequencing for De Novo Assembly

We selected 20 dihaploids based on their greenhouse phenotypes and sequence diversity to generate *de novo* assemblies. For these individuals, high molecular weight DNA was extracted from above ground tissue from seedlings grown in tissue culture using a CTAB isolation method in combination with a Qiagen Genomic Tip (Hilden, Germany) followed by an Amicon filter (MilliporeSigma, Burlington, MA) buffer exchange (Vaillancourt and Buell, 2019) or Takara NucleoBond HMW DNA kit (Takara, Kusatsu, Shiga, Japan). PacBio HiFi libraries were prepared and sequenced by the University of Minnesota Genomics Center. First, high molecular weight DNA was Pippin size-selected as needed and libraries prepared using either the SMRTbell Express Template Prep Kit 2.0 or SMRTbell Template Prep Kit 3.0 (Menlo Park, CA). All sequencing was performed on a PacBio Sequel II with an output of 31.4 - 56.8X coverage per haplotype (Supplemental Table 12).

PacBio HiFi reads were input into hifiasm v0.16.1-r375 (Cheng et al. 2021, 2022) to generate haplotype-resolved assemblies. To account for the variability in heterozygosity across the set of 20 dihaploids, hifiasm was run with a similarity threshold (-s) of .2, .35, .4, .45, and .5 for each dihaploid. The total number of contigs, assembly size, and N50, were calculated using assembly-stats (https://github.com/sanger-pathogens/assembly-stats) and assembly completeness was evaluated using BUSCO v5.4.3 (Manni et al. 2021) with the embryophyta_odb10 lineage database. These results were used to determine the similarity threshold to proceed with for each individual dihaploid. Contigs less than 50 kb were discarded using seqkit v2.3.0 (Shen et al. 2016). Two rounds of Ragtag v2.1.0 (Alonge et al. 2019) were then run to scaffold the two individual haplotypes with DM 1-3 516 R44 v6.1 (Pham et al. 2020) as the query genome. To check for contamination both the PacBio HiFi reads and assembly were split into 150 bp fragments using seqkit v2.3.0 0 (Shen et al. 2016) and then Kraken v2.1.2 (Wood et al. 2019) was run using the k2_pluspfp_20220908 database (PlusPFP includes Refeq bacteria, viral, plasmid, human, UniVec Core, protozoa, fungi, and plant) to check for contamination; unclassified, viridiplantae, bacteria, archaea, and virus percentages were evaluated and no contamination was found in the datasets. Each haplotype assembly was then aligned to the DM 1-3 516 R44 v6.1 (Pham et al., 2020) genome using minimap2 (Li, 2021) with the parameters -x asm10 and -secondary=no; alignments were then plotted using D-Genies (Cabanettes and Klopp 2018) to determine chromosomes that were unphased (Supplemental Figure 2).

### Genome Annotation

To provide empirical evidence for gene annotation, total RNA was isolated from immature leaf and whole tuber tissue using a hot borate method (Wan and Wilkins 1994) and DNA removed using Invitrogen TURBO DNase (Waltham, MA). mRNA was purified from total RNA using the Invitrogen Dynabeads mRNA Purification Kit (Waltham, MA) and then libraries made using Oxford Nanopore Technologies (ONT) PCR-cDNA Barcoding Kit SQK-PCB109 (Oxford, United Kingdom). All libraries were sequenced on FLO-MIN106 Rev D flow cells via a MinION sequencing device (Supplemental Table 13). Base calling was performed using Guppy v6.1.7 (Kalkaska-DH031), v6.2.1 (A07061-6-DH007, Missaukee-DH157, PayetteRusset-DH004, and W14NYQ29-5-DH024) or v6.5.7 (rest of libraries) with the following parameters: -- config dna_r9.4.1_450bps_sup.cfg -q 0 --trim_strategy none --barcode_kits "SQK-PCB109" -- calib_detect.

Dihaploid genome assemblies were annotated for protein-coding genes as described previously (Pham et al. 2020). In brief, the genome assemblies were repeat masked by first creating a custom repeat library (CRL) for each genome. Repeats were first identified with RepeatModeler (v2.03; Flynn et al. 2020) and protein coding genes filtered out from the repeat database using ProtExcluder (v1.2; Campbell et al. 2014) to create a CRL. The CRL was then combined with Viridiplantae repeats from RepBase (v20150807; Bao et al. 2015) to generate the final CRL for each genome. Each genome assembly was repeat-masked using the respective final CRL and RepeatMasker (v4.1.2-p1; Chen 2004) using the parameters -e ncbi -s -nolow -no_is -gff.

Oxford Nanopore (ONT) cDNA reads were processed with Pychopper (v2.5.0; github.com/nanoporetech/pychopper) and trimmed reads greater than 500 nt were aligned to the respective genome using minimap2 (v2.17-r941; Li, 2018) with a maximum intron length of 5,000 nt. The aligned ONT cDNA reads were each assembled using Stringtie (v2.2.1; Kovaka et al., 2019) and transcripts less then 500 nt were removed.

The initial gene models for each genome were created using BRAKER2 (v2.1.6; Hoff et al. 2019) using the soft-masked genome assemblies and the Atlantic potato training matrix (Hoopes et al. 2022). The gene models were then refined using the ONT transcript assemblies with two rounds of PASA2 (v2.5.2; Haas et al. 2003; Campbell et al. 2006) to create a working gene model set for each genome. High-confidence gene models were identified from each working gene model by filtering out gene models without expression evidence, a PFAM domain match, a partial gene model, or contained an interior stop codon. Functional annotation was assigned by searching the working gene models proteins against the TAIR (v10; Lamesch et al. 2012) database and the Swiss-Prot plant proteins (release 2015_08) database using BLASTP (v2.12.0; Altschul et al. 1990) and the PFAM (v35.0, El-Gebali et al. 2019) database using PfamScan (v1.6) and assigning the annotation based on the first significant hit.

### GENESPACE Syntelog Analysis

Syntelogs were identified between the 20 dihaploid potato genomes, DM potato (v6.1; Pham et al. 2020), Atlantic (v3; Hoopes et al. 2022), Castle Russet (v2; Hoopes et al. 2022), Otava (Sun et al. 2022), and Cooperation-88 (Bao et al. 2023) using GENESPACE (v1.2.3; Lovell et al. 2022) with the representative working model proteins for each genome and setting the expected ploidy. An additional GENESPACE run was performed with the haplotypes of the polyploid genomes separated and the ploidy set to 1 for each haploid genome or haplotype. The syntelog sets were extracted from the pangenome database file from each GENESPACE run.

## Data Availability

Raw sequence data is available at the National Center for Biotechnology Sequence Read Archive under BioProject PRJNA1055108 (data to be released upon publication) and assemblies via SpudDB (spuddb.uga.edu) and Figshare (https://doi.org/10.6084/m9.figshare.29803324.v1).

## Acknowledgements

We acknowledge the technical assistance of Lemor Carlton, Rachel Shereda, and Joshua C. Wood. Financial support was provided by the USDA National Institute of Food and Agriculture, Award 2019-51181-30021, Pepsi Co, and the Minnesota Department of Agriculture through AGREETT. We acknowledge financial support from pre-doctoral Netaji Subhas ICAR International fellowship, Education Division, Indian Council of Agriculture Research, Krishi Anusandhan Bhavan II, Pusa, New Delhi-110012 for Hemant Balasaheb Kardile. We thank Dr. Ya Yang for her comments on the draft.

